# Histological and haematological analysis of the ascidian *Botrylloides leachii* (Savigny, 1816) during whole-body regeneration

**DOI:** 10.1101/099580

**Authors:** Simon Blanchoud, Lisa Zondag, Miles D. Lamare, Megan J. Wilson

**Affiliations:** Department of Anatomy, Otago School of Medical Sciences; University of Otago, P.O. Box 56, Dunedin 9054, New Zealand.; Present address: Department of Biology, University of Fribourg, Chemin du Musée 10, 1700 Fribourg, Switzerland.; Department of Marine Science, Division of Sciences; University of Otago, P.O. Box 56, Dunedin 9054, New Zealand.

**Keywords:** *Botrylloides leachii*, haemocytes, haemolymph flow, histology, whole-body regeneration

## Abstract

Whole-body regeneration, the formation of an entire adult from only a small fragment of its own tissue, is extremely rare among chordates. Exceptionally, in the colonial ascidian *Botrylloides leachii*, a fully functional adult is formed from their common vascular system, upon ablation of all adults from the colony, in just 10 days thanks to their high blastogenetic potential. While previous studies have identified key genetic markers and morphological changes, no study has yet focused on the haematological aspects of regeneration despite the major involvement of the remaining vascular system and the contained haemocytes in this process. To dissect this process, we analysed colony blood flow patterns using time-lapse microscopy to obtain a quantitative description of the velocity, reversal pattern, and average distance travelled by haemocytes. We also observed that flows present during regeneration are powered by temporally and spatially synchronized contractions of the terminal ampullae. In addition, we revised previous studies on *B*. *leachii* haematology as well as asexual development using histological sectioning, and compared the role of haemocytes during whole-body regeneration. We found that regeneration starts with a rapid healing response characterized by blood clotting and infiltration of immunocytes, followed by increased activity of haemoblasts, recruitment of macrophage-like cells for clearing the tissues of debris, and their subsequent disappearance from the circulation concomitant with the maturation of a single regenerated adult. Overall, we provide a uniquely detailed account of the haematological properties of regenerating *B*. *leachii* colonies, providing novel lines of inquiry towards the decipherment of regeneration in chordates.

## Introduction

Whole-body regeneration (WBR) is the process whereby an entire functional adult is formed from only a minute fragment of the original organism. Multicellular animals capable of varying degrees of regeneration are distributed widely throughout the metazoa (Sánchez Alvarado and Tsonis, 2006), thus suggesting that this biological phenomenon has a primordial origin. However, this regeneration ability correlates inversely with body and tissue complexity (Alvarado, 2000; Rinkevich et al., 2007).

Among vertebrates, while most adults heal only through scarring, the teleost fish and urodele amphibians can regenerate tissues, body parts or organs following injury (Jaźwińska and Sallin, 2016; Seifert et al., 2012; Tanaka and Reddien, 2011). While the study of these organisms has brought an extensive knowledge of patterning and cellular programs required for regeneration in mature animals following severe injury, no vertebrate has yet been shown to undergo WBR. Therefore, deciphering the mechanisms underlying regeneration in evolutionarily related organisms might provide new implications about the basic principles of cellular plasticity in adult organisms of our phylum.

Colonial tunicates represent the closest phylogenetic relative of vertebrates (Delsuc et al., 2006) and can undergo WBR due to their high blastogenetic potential (Oka & Watanabe, 1957, 1959; Milkman, 1967; Zaniolo et al., 1976; Rinkevich et al., 1995). Belonging to the Phylum Chordata, Class Ascidiacea, these invertebrates have tissue and organ complexity reminiscent of vertebrates (see Fig. 1), including a well-developed notochord, dorsal nerve cord, and a post-anal tail during their free-living tadpole larval stage. In particular, *Botrylloides leachii* (order Stolidobranchia, family Styelidae; Savigny, 1816) can undergo WBR in as little as 10 days (Rinkevich et al., 1995; Rinkevich et al., 2007, 2008, 2010; Zondag et al., 2016). Consequently, *B*. *leachii* has recently emerged as a model organism for the study of regeneration (Gross, 2007).

**Fig. 1.**
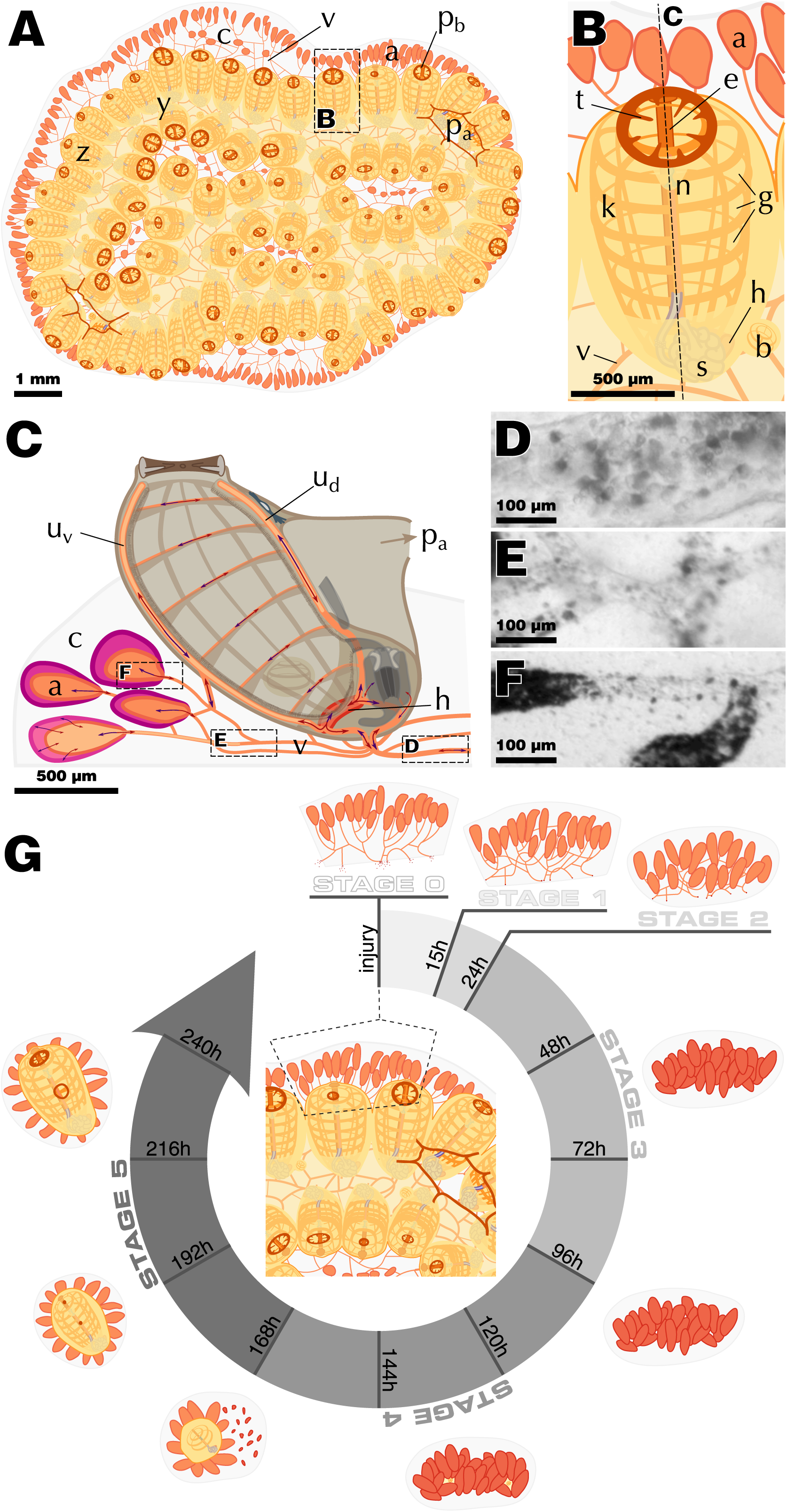
*Botrylloides leachii* schematic morphology and physiology. Abbreviations: zooid (z), system (y), tunic (c), vascular system (v), terminal ampullae (a), buccal siphon (pi,), atrial siphon (p_a_), oral tentacles (t), endostyle (e), pharyngeal basket (k), stigmata (g), neural ganglion (n), heart (h), stomach (s), bud (b), dorsal sinus (ud), ventral sinus (u_v_). (**A**) Top-view of a stereotypical colony composed of 72 zooids. Dashed-delimited area magnified in **B**. (**B**) Top-view of a single zooid. Longitudinal plane (dashed) depicted in **C**. (**C**) Left lateral view of a zooid. Abvisceral/advisceral haemolymph direction and contraction/dilatation of ampullae are depicted using red/purple arrows, respectively. Dashed-delimited areas are exemplified in **D-F**. *In vivo* stereo-microscopic images of a large vessel (**D**), vascular junction (**E**) and terminal ampullae (**F**), taken from recordings showcased in Movie SI. (**G**) *B*. *leachii* WBR from a fragment of vascular tissue through our five defined stages and the approximate time-line

*B*. *leachii* are sessile suspension-feeding ascidians that live in clonal colonies of adults, termed zooids, which organize in a series of ladder-like parallel rows, known as systems (Fig. 1 A-B; Brewin, 1946; Michaelsen, 1921; Savigny, 1816). The colony attaches to its substrate, typically underneath rocks in the shallow subtidal zone, through a gelatinous tunic. While each zooid has an independent heart and an open haemocoelic circulatory system, the entire colony shares a common vascular system that is embedded in the tunic and closed at its periphery by terminal ampullae, the contractile blind ends of marginal vessels (Fig. 1 C-F; Mukai et al., 1978). *B*. *leachii* zooids are hermaphrodites and can reproduce both sexually, to colonize new locations through a tadpole larval stage, or asexually, to expand the colony (Berrill, 1947; Mukai et al., 1987). This latter reproduction, also known as palleal budding (Oka and Watanabe, 1959) or blastogenesis (Brunetti, 1976), occurs on the outer epithelial mantle of a zooid, typically on both of its sides thus producing two offspring. These growing buds start as a thickened ring of cells on the body wall and invaginate to form the required layers of tissue for organ development. Eventually, these buds, located on either side of the older zooid, will mature into new adults (Berrill, 1947). As in the closely related genus *Botryllus* (Berrill, 1941; Sabbadin, 1969), bud development is synchronous throughout the colony and starts very early in the life cycle of the parent bud (Berrill, 1947; Brunetti, 1976).

In addition to these two methods of reproduction, *B*. *leachii* colonies can undergo WBR from a minute fragment of their vascular tissue (∼200 cells; Rinkevich et al., 1995). WBR will only be triggered following the loss of all zooids from the colony; otherwise a more traditional injury healing will occur (Rinkevich et al., 1995). *Botrylloides leachii* WBR starts with the healing of the injury sites to prevent further haemolymph loss, followed by the compaction of the marginal ampullae towards the centre of the remaining matrix. Within this reorganized tissue, regeneration niches - a discrete regeneration locus within the vascular system - will develop and compete through a yet undetermined process that ultimately leads to the development of a single adult zooid, while all other niches are absorbed by the colony (Fig. 1 G). Importantly, WBR ability in *B*. *leachii* is retained throughout its developmental cycle (Oka and Watanabe, 1959; Rinkevich et al., 2007), in contrast to the sister species *Botryllus schlosseri*, where WBR potential is only present during a one-day period of their asexual reproduction cycle, known as takeover (Rinkevich et al., 2008; Voskoboynik et al., 2007). While the exact origin of the totipotent stem-like cells responsible for WBR remains unknown, it has been shown that even though they will develop a regeneration niche within the vascular system, they are not part of the circulating blood cells (haemocytes) in uninjured colonies but rather appear to migrate from the vascular lining into the vascular system after injury (Rinkevich et al., 2010).

Similar to most other chordates, ascidian haemocytes are continually renewed to compensate for cell death and external sources of loss (Ermak, 1982). The location of haemopoiesis in ascidians is still debated, which might be a consequence of the variety of developmental strategies observed among this clade (Brown and Swalla, 2012). In some solitary ascidians, it principally occurs in small static clusters of haemoblasts located in the lining of the haemocoel, particularly around the pharyngeal basket (Ermak, 1982). In colonial ascidians, clusters of undifferentiated cells with pluripotent properties, called islands, have been detected in the sub-endostylar ventral sinus (Voskoboynik et al., 2008). In addition, ventral islands located near the endostyle have been identified as sites of phagocyte turnover (Lauzon et al., 2013). Furthermore, circulating haemoblasts have been shown to express stemness and proliferation markers around early buds of *Botrylloides violaceus*, suggesting a circulatory source of haemocytes (Brown et al., 2009). It thus appears that there may be multiple sites for ascidian blood (haemolymph) production, in particular in colonial tunicates.

Haemolymph is mostly colourless and composed of a variety of cell types with functions reminiscent to that of vertebrates (Cima et al., 2002; Goodbody, 1975; Millar and Ratcliffe, 1989; Wright, 1981); and its flow patterns are intimately linked to the health of the whole colony (Dijkstra et al., 2008).

Although there has been a number of studies investigating the morphotypes of ascidian haemocytes (Ballarin et al., 2011; Ballarin and Cima, 2005; de Leo, 1992; Endean, 1960; Freeman, 1964; Hirose et al., 2003; Wright and Ermak, 1982), including offi *leachii’s* (Cima et al., 2002), improved resolution and colour images would benefit future work on this species. In addition, no common terminology has been accepted to identify these cell types, mainly because of discrepancies between their phenotype and their function in different species, as well as a lack of understanding of their differentiation lineages (Ballarin et al., 2011; Cima et al., 2002; Endean, 1960; Freeman, 1964; Goodbody, 1975; Millar and Ratcliffe, 1989; Wright, 1981). Furthermore, accurate identification is crucial for detecting and interpreting variations in the sub-populations of haemocytes during WBR.

Despite the major involvement of the remaining vascular system and the comprised haemocytes during WBR in *B*. *leachii*, no study has yet focused on the haematological aspects of regeneration. To obtain a more accurate understanding of the WBR process in *B*. *leachii*, we set out to characterize in a consistent and amended approach both uninjured and regenerating colonies at the circulatory, tissue, and cellular levels. We used both histological staining and time-lapse recording to dissect this highly dynamic process of regeneration with greater resolution.

## Materials and Methods

### Animal husbandry and manipulation

*Botrylloides leachii* colonies were collected from the two sites in the Otago Harbour (45°52’12”S, 170°31’48”E and 45°49’41”S, 170°38’29”E), and Nelson Harbour (41°15’36”S, 173°16’48”E), New Zealand between September and March over 2014 to 2016. *B*. *leachii* grows naturally on submerged structures (e.g. ropes, pontoons, tyres), and colonies were removed from the attached substrata with a single edge razor blade. Each colony was placed on either 5.0 × 7.5-cm or 2.6 × 7.6-cm glass slides and left horizontally for two days in 200 ml of still seawater, changed daily, to allow the colony to attach to the slide. The slides were then placed vertically and kept at the Portobello Marine Laboratory (University of Otago) in a tank supplied with a constant flow-through of filtered seawater directly from the harbour, with temperature controlled between 18 °C and 20 °C. The glass slides were kept clear of algal growth by using a paintbrush and a single edge razor blade (after Rinkevich et al., 2007). Colonies were continuously fed, using a peristaltic pump, a 1:1:1:1 by volume mixture of three algae (*Chaetoceros muelleri*, *Pavlova lutheri* and *Tetraselmis chuii*) and a rotifer (*Brachionus plicatilis*) culture, complemented with Alltech All-G-Rich algae/yeast powder mix (12 g of Alltech All-G-Rich were added each week to the rotifer culture).

*B*. *leachii* haemolymph was collected as previously described (Cima, 2010). Briefly, whole colonies were first cleaned using filtered seawater (FSW) and incubated for 5 min in anticoagulant solution (0.38 g sodium citrate per 100 ml FSW, pH 7.5). To prevent loss of haemocytes, all instruments were incubated twice for 1 min in the anticoagulant solution prior to collection. Colonies were then dried using blotting paper and their marginal vessels broken using a needle. The haemolymph was then collected using a syringe and transferred into a 1.5-ml microfuge tube containing 100 μl of the anticoagulant solution up to a total volume of 600 μl. The tube was left 5 min on ice and the supernatant transferred to a new tube to remove debris. Following centrifugation at 780 g for 12 min, the supernatant was discarded and the pellet re-suspended in 100 μl FSW.

*B.leachii* regeneration was carried out following an established protocol (Rinkevich et al., 2007). Dissection of adults and buds away from the marginal ampullae, during their midcycle blastogénie stage (i.e. not undergoing take-over), was carried out using a scalpel and a single edge razor blade. The slides with the remaining vascular fragments were then placed into an aerated seawater tank at 19 °C and left to regenerate until a certain morphological stage had been reached.

### Histological staining

*B. leachii* regenerating tissue fragments were fixed overnight in 4% paraformaldehyde (PFA) in FSW, dehydrated in 70 % ethanol (EtOH) and embedded in paraffin wax. The fragments were then sectioned (5 μm thicknessj in the transverse plane. Three stains were used on dewaxed slides: haematoxylin and eosin (H&E), Giemsa, and Martius Scarlet Blue trichrome (MSB). H&E staining was performed as follows: stain for 4 min in haematoxylin (Leica Biosystems Surgipath Gill II Haematoxylin), wash for 2 min in running tap water, place the slide for 2 min in Scott’s tap water (2 g of potassium bicarbonate, 20 g magnesium sulphate per litre), wash again for 2 min, stain for 30 s in eosin (Leica Biosystems Surgipath Eosin), wash for 1 min, dehydrate and mount Giemsa staining was performed as follows: stain for 30 min in a coplin jar containing Giemsa (7.36 g Giemsa powder in 500 ml of 50 °C glycerol and 500 ml methanol, mix 1:45 in dH_2_O) within a 56 °C waterbath, rinse in dH_2_O, differentiate in 1/1500 acetic acid solution, rinse in dH_2_O, dehydrate and mount. MSB staining was performed as follows: stain for 5 min in Celestine blue (5 g ferric ammonium sulphate, 0.5 g Celestine Blue B in 100 ml dH_2_0 with 14 ml glycerol), wash for 30 s in running tap water, stain for 5 min in haematoxylin, wash again for 2 min, place the slide for 2 min in Scott’s tap water, wash for 2min, rinse for 1min in 95 % ethanol, stain for 2 min in martius yellow (20 g phosphotungstic acid, 5 g martius yellow per litre of 95 % ethanol), rinse three times for 1 min in dH_2_O, stain for 10 min in brilliant scarlet (10 g brilliant scarlet crystal ponceau 6R, 20 ml glacial acetic acid per litre of dH_2_O), rinse again three times, place the slide for 4 min in phosphotungstic acid (10 g per litre of dH_2_0), rinse for 1 min in dH_2_O, stain for 45 s in methyl blue (5 g methyl blue, 10 ml glacial acetic acid per litre of dH_2_O), rinse for 1 min in acetic acid (10 ml glacial acetic acid per litre of dH_2_O), dehydrate and mount

Haemolymph smears were obtained as previously described (Cima, 2010). Briefly, drops of 50 μm of isolated haemocytes (see above) were left for 30 min to attach on SuperFrost Plus (Thermo Fisher Scientific) microscopy slides. The drop was then discarded by placing the slide vertically and the remaining attached cells were fixed for 30 min at 4 °C in ascidian fixative solution (1 g NaCl and 1 g sucrose in 1 % glutaraldehyde in FSW). Slides were then rinsed for 10 min in 0.1 M PBS, stained using 10 % Giemsa, mounted using a 9:1 glycerol-PBS solution and sealed with a coverslip using nail polish.

### Imaging

All fixed samples were imaged on an Olympus AX70 microscope equipped with a QImaging MicroPublisher 5.0 2560 x 1920 pixels camera using QCapture. Magnification used ranged from 4 x to 100 x. Time-lapse recordings of haemolymph flow in live colonies were acquired on a Leica M205 FA stereo-microscope equipped with an 8 megapixel CCD DFC490 colour camera using the Leica Application Suite (LAS). Recordings were acquired using the “Movie” continuous mode, at different magnifications (7.42 x to 20 x) and exposure times (27.7 ms to 55.7 ms) adapted to the recorded portions of the colony, with a resolution of either 544 x 408 or 1088 x 816 pixels.

### Data processing

Acquired images were processed for colour balance using the open-source Gimp program (www.gimp.org) and assembled into figures using the open-source Inkscape software (www.inkscape.org). Colours were chosen according to the ColorBrewer palette (www.colorbrewer2.org). Movies were montaged using the Lightworks program (www.lwks.com) and compressed using the open-source HandBreak tool (www.handbreak.fr).

Haemolymph composition during WBR was estimated by manually identifying cell types in microscopy images using five randomly located images of Giemsa stained ampullae, at a 60-x magnification. Similarly, 10 images were used to characterize haemolymph composition in intact colonies fixed during their mid-cycle blastogénie stage. Only cells located inside the ampullae, centred on the imaged section and unambiguously identified (2.6 % of the total 2339 cells could not be determined) were considered. Between 272 and 447 cells were positively identified for each stage of regeneration, 502 for the intact colony. We focused our counting on ampullae for setting up a meaningful comparison between intact and regenerating tissues as blood vessels become very sparse after tissue contraction (i.e. stage 3, see Fig. 5).

The appearance of haemocytes was characterized using light microscopy by smearing ascidian haemolymph and by staining histological sections of whole colonies.

### Computational analysis

Vessel identification in time-lapse recordings was performed as follows. Raw frames (Fig. S1 A) of the recording were smoothed using a Gaussian kernel of radius 0.89 μm the difference between consecutive frames was computed (Fig. S1 B) and moving cells were identified as having an absolute value greater than 6 x the estimated standard deviation of the image’s white noise (Fig. S1 C; Paul et al., 2010). Cells larger than 5 μm^2^ were then consecutively morphologically dilated and eroded, using structuring disks of radii 41 μηι and 21 μηι, respectively. The resulting detections were then averaged over the entire recording (Fig. S1 D), and vessels defined as containing moving cells in at least 30 % of the frames (Fig. S1 E). The location, length and width the vascular systems were then identified using the morphological skeleton of the vessels (Fig. S1 F).

Haemolymph flow was measured using a particle image velocimetry (PIV) approach with multiple passes and subpixel resolution using Fourier-space (Liao and Cowen, 2005) based on the difference between consecutive frames. Raw frames of the recording were smoothed using a Gaussian kernel of radius 0.89 μηι, the difference between consecutive frames was computed (Fig. S1 G-I) and PIV performed between two consecutive differences, using a modified version of matpiv_nfft from the MatPIV toolbox (http://folk.uio.no/jks/matpiv/index2.html), through four consecutive passes with interrogation windows of sizes 83 x 83, 62 x 62, 41 x 41 and 21 x 21 μm; and finalized with a subpixel pass in Fourier-space (Fig. S1 J & K). Only the speeds measured inside the previously detected vessel segments were retained and projected onto the tangent of the segment. The orientation of each segment was then adjusted based on the maximal correlation between average speeds. Speeds from all aligned segments were then binned into a 2D histogram of resolution 350 ms x 6 μm/s, and the most likely haemolymph flow was computed using a shortest path approach (Dijkstra, 1959), implemented using dynamic programming (Fig. S1 L; Bellman, 1952). Finally, to dampen the effect of the binning required by dynamic programming, the resulting haemolymph flow was smoothed using cubic smoothing splines (Fig. S1 L).

Ampullar contractions were measured as follows. We first identified ampullae in every frame (Fig. S2 A) as regions darker than 20 x the estimated standard deviation of the image’s white noise (Paul et al., 2010). The resulting binarized image was then consecutively morphologically opened and closed, using a structuring disk of radius 9.5 μm (Fig. S2 B). The area and the centroid of each separated object was computed (Fig. S2 C), and the trajectory of each ampulla was reconstructed by clustering across frames detections whose centroids were closer than 19 μm (Fig. S2 D). Only trajectories that spanned more than 95 % of the frames were kept for further analysis. All the code written for our analysis was implemented as a set of custom MATLAB R2015b (MathWorks) functions and is available upon request

## Results

### Haemolymph circulation

As a first step towards dissecting WBR, we characterized the haematological properties of a healthy and intact adult colony of *B*. *leachii* (Fig. 1 A-B). Haemolymph circulation in suspension-feeding ascidians undergoes a periodic reversal of flow direction between advisceral (towards the viscera) and abvisceral (towards the branchial basket). The advisceral flow starts on the anterior side of the heart, moves towards the endostyle before arriving to the pharynx and the mid-dorsal vessel, which supplies haemolymph to the digestive tract, and lastly passes to the dorsal end of the heart (Mukai et al., 1978).

Utilising the transparency of the tunic, we recorded time-lapse stereomicroscopy data from eight colonies at different locations in their vascular system (Fig. 1 C-F, Movies S1). Focusing on the largest vessels of the colony (Movie S1-S2), we gathered a set of reproducible flow measurements (n=9, from 5 different colonies) that allowed us to calculate, using a custom-made PIV-based tracking software, the flow (peak velocity ˜200μm/5) and reversal rates (∼57/24 s) of a *B*. *leachii* colony (Fig. 2 A & Fig. S1).

**Fig. 2.**
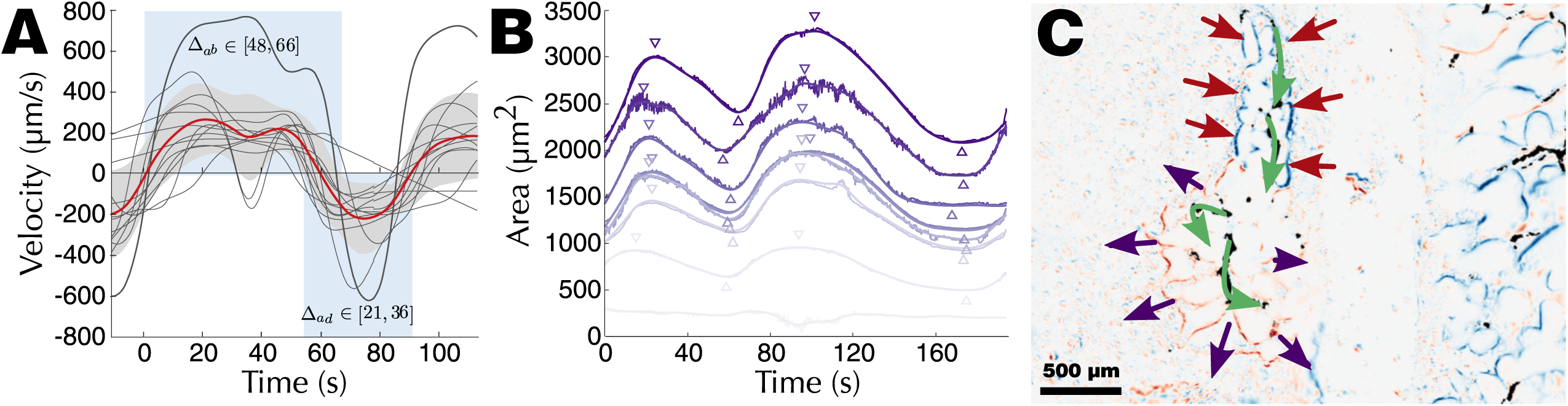
Quantification of the haemolymph circulation in *B. leachii.* (**A**) Average (n=8) abvisceral haemolymph flow based on PIV measurement of time-lapse large vessel recordings. Depicted are the individual recordings (thin grey lines), the recording utilized for Fig. S1 (thick grey line), the average velocity (red line) and standard deviation (light grey area). A visual representation of the duration of abvisceral and advisceral phase durations is shown (blue). (**B**) Dilatation-contraction cycles of eight terminal ampullae (Fig. S2) from a single time-lapse recording. The raw data (darker line) is overlaid with a smoothed measurement (lighter line), local maxima and local minima (down and up triangles, respectively). The abvisceral phase of the haemolymph flow corresponds to the contraction phase of the ampullae. (**C**) Ampullae contractions are independent from heart activity and can drive haemolymph flow (green arrows) by synchronized spatially restricted contractions (red arrows) and dilatations (purple arrows)

When recording vascular junctions, we observed (Movie S1) and quantified (Movie S3) an apparently pseudo-erratic pattern at the time of flow reversal. By examining terminal vessels (Movie S1), we extracted valuable information on the rate (11.4 ± 5.3 μm^2^/s, n=18) and extent (dilated/contracted surface ratio: 1.55 ± 0.26, n=12) of the ampullar contraction-dilatation cycle concomitant with haemolymph flow in ascidians (Mukai et al., 1978). While the general timing of haemolymph and ampullae alternation is synchronized (average lag: 2.9 ± 2.5 s, n=24), small variations on the exact time of contraction for each ampulla could be measured (± 3.1 s, n=24,_Fig. 2 B &Fig. S2).

### Haemocyte classification

By combining previous classification schemes for botryllid ascidians (Ballarin et al., 2011; Ballarin and Cima, 2005; Cima et al., 2002; Hirose et al., 2003), we propose a generalized classification of *B*. *leachii* circulatory haemocytes into five functional groups: undifferentiated cells, immunocytes, mast cell-like cells, transport cells and storage cells (Table 1). A further detailed description of the observed cell types is provided in Table SI. In this classification, undifferentiated cells are composed solely of haemoblasts, also called lymphocytes or lymphocyte-like cells. Immunocytes can be further classified between phagocytes and cytotoxic cells. Phagocytes include hyaline amoebocytes and macrophage-like cells while cytotoxic cells include granular amoebocytes and morula cells. Mast-like cells are represented only by granular cells. Transport cells are composed of compartment amoebocytes and compartment cells. Finally, storage cells include both pigment cells and nephrocytes. A comprehensive guide to haemocyte identification using bright field Giemsa stained images is provided (Table 2), with high magnification pictures exemplifying the most common aspect of each cell type (Fig. 3).

**Table 1.** Summary of cell types present in the *Botrylloides leachii* circulatory system.

| Cell type | Subtype | Circulatory cells | Older terminology |
| --- | --- | --- | --- |
| Undifferentiated cell <sup>c, d</sup> | Haemoblast <sup>e, f</sup> | Haemoblast <sup>e, f</sup><br>Differentiating cell | Lymphocyte <sup>a, b</sup><br>Stem cell <sup>d</sup> |
| Immunocyte <sup>c</sup> | Phagocyte <sup>e</sup> | Hyaline amoebocyte <sup>d, f</sup><br>Macrophage-like cell <sup>d, f</sup> | Amoebocytes with vacuole<br>Signet ring cell <sup>a, b</sup><br>Vacuolated cell <sup>c</sup><br>Leukocyte <sup>c</sup> |
|  | Cytotoxic cell <sup>f</sup> | Granular amoebocyte <sup>b, d, f</sup><br>Morula cell <sup>b, d, e</sup> | Cells with acidic vacuole <sup>a</sup><br>Green cell <sup>b</sup><br>Leukocyte <sup>c</sup><br>Vanadocyte <sup>a</sup> |
| Transport cell <sup>e</sup> | Granulocyte <sup>e</sup> | Compartment amoebocyte <sup>d</sup><br>Compartment cell <sup>a, b, d</sup> | Vacuolated cell <sup>c</sup> |
| Mast cell-like cell <sup>d</sup> | Granular cell <sup>d</sup> | Granular cell <sup>d</sup> | (Unique to *B. leachii*, <sup>d</sup>) |
| Storage cell <sup>f</sup> | Vacuolated cell <sup>f</sup> | Pigment cell <sup>a, c, d, e</sup> | Cell with reflecting disks <sup>a</sup> |
|  |  | Nephrocyte <sup>c, d, e</sup> | Orange cell <sup>b</sup> |

<sup>a</sup> (Endean, 1960); <sup>b</sup> (Freeman, 1964); <sup>c</sup> (Wright & Ermak, 1982); <sup>d</sup> (Cima et al., 2002); <sup>e</sup> (Hirose et al., 2003); <sup>f</sup> (Ballarin et al., 2011)

**Fig. 3.**
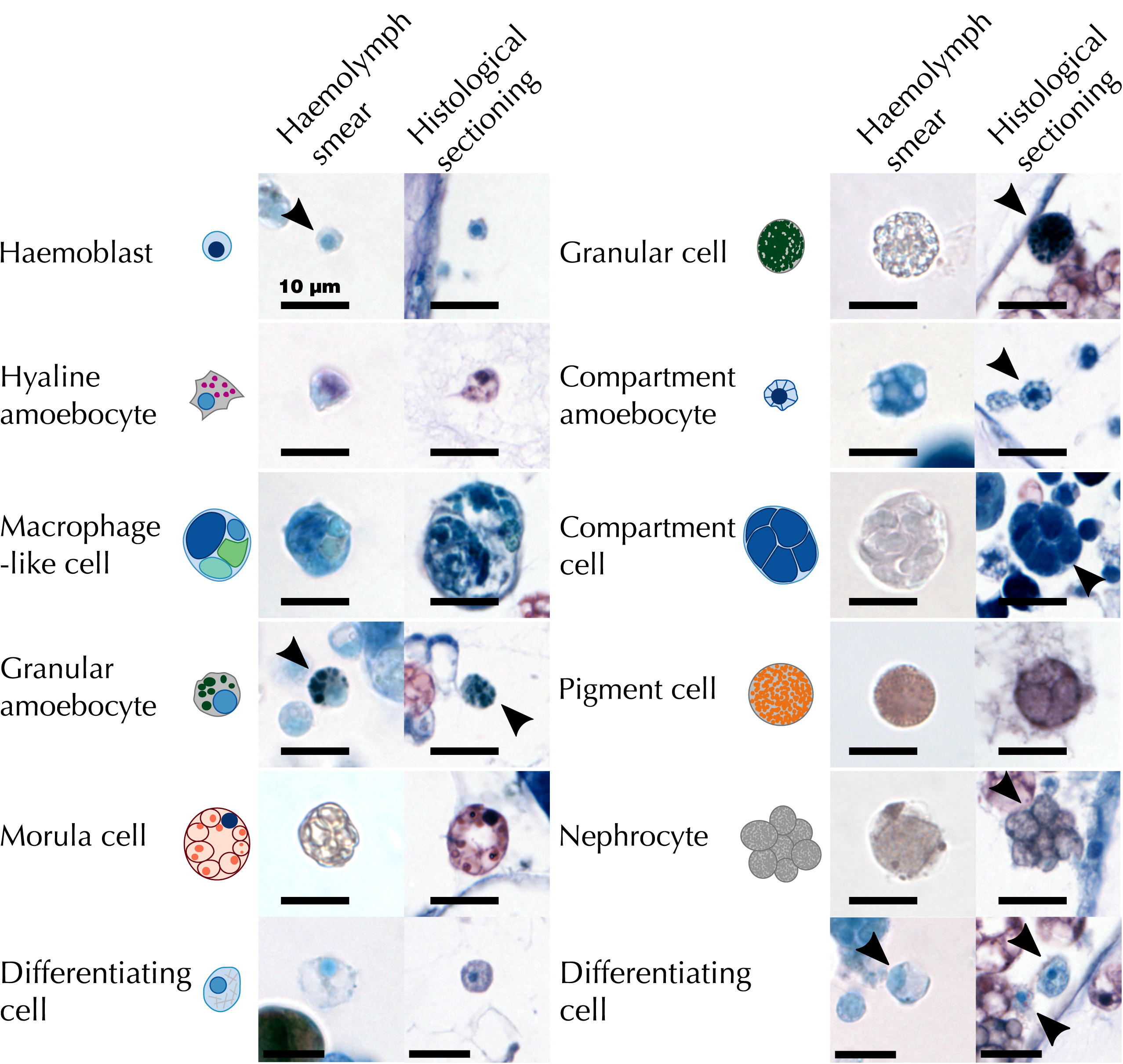
Characterization of *B*. *leachii* haemocytes. Stereotypical schemas along with characteristic images of all haemocytes identified in *B*. *leachii* stained with Giemsa dye, both from haemolymph smears and whole-colony histological sections. The illustrated cell is either located at the centre of the image or indicated by an arrowhead. Note that the cells in the histological sectioning show a degree of shrinkage following PFA fixation

Altogether, these descriptions (Table 2) and images (Fig. 3) allowed us to identify most of the cells both in haemolymph smears and histological sections and provide an unambiguous identification reference for the analysis of the localization and fate of *B*. *leachii* haemocytes in the colony.

**Table 2.** Classification chart of *Botrylloids leachii* haemocytes based on light microscopy Giemsa stain. Only the most common traits are listed in this table, refer to Table SI for full description of the cell types.?

| Cell type | Size (µm) | Shape | Stain | Characteristic |
| --- | --- | --- | --- | --- |
| Haemoblast | 5 - 8 | Round | Blue cytoplasm with a dark-blue nucleus | High nucleus to cytoplasm ratio |
| Differentiating cell | 8 - 12 | Swollen, irregular | Light-blue nucleus and grey cytoplasm | Fibrous cytoplasm with a proportionally large nucleus |
| Hyaline amoebocyte | 6 - 9 | Amoebocyte: highly irregular, including a variety of pseudopodia | Dual purple-blue cytoplasm | Purple stained clear cytoplasmic vesicles |
| Macrophage-like cell | 6 - 20 | Mostly round with irregular content | Blue with vesicles of varying shades of yellow, brown, green and grey | Vesicles highly heterogeneous in size, shape and content |
| Granular amoebocyte | 6 - 10 | Amoebocyte | Scattered dark-green cytoplasmic vesicles | Sparser vesicles than granular cells |
| Morula cell | 10 - 16 | Circular containing a low number of large (~2 µm), pale yellow and round vacuoles | Orange-green cytoplasmic colour in smears, red colour around the vacuoles in histological sections | Most common cell type in *B. leachii* haemolymph |
| Granular cell | 8 - 12 | Circular with irregular edge | Dense dark-blue/green vesicles | High density of vesicles renders the cell black and opaque |
| Compartment amoebocyte | 7 - 10 | Amoebocyte with a berry-like shape | Blue with a dark-blue nucleus | Numerous clear protruding vesicles |
| Compartment cell | 12 - 18 | Similar to morula | Unstained-grey opaque vesicles | Large uniform vesicles |
| Pigment cell | 10 - 16 | Similar to morula | Filled with minute dark granules, related to the colony colour | The content of the vesicle exhibits Brownian movement in living cells |
| Nephrocyte | 10 - 16 | Berry-like | Dark and speckled grey vesicles | An aggregate of dark grey vesicles |

### Asexual budding

To obtain a precise picture of the prevailing mode of asexual reproduction, we revised various stages of blastogenesis in *B*. *leachii* previously described by Berrill (1941, 1947) using histological sections. Palleal reproduction involves the formation of a new bud from the peribranchial epithelium of a zooid. Remarkably, in *B*. *leachii* as well as in other colonial botryllid, because blastogenesis starts very early in the development of a maturing bud, there are typically three generations co-existing in the colony (Fig. 4 A): second-generation buds (budlets), developing from more differentiated buds (first-generation buds), themselves attached to adult filter-feeding zooids (blastozooids).

**Fig. 4.**
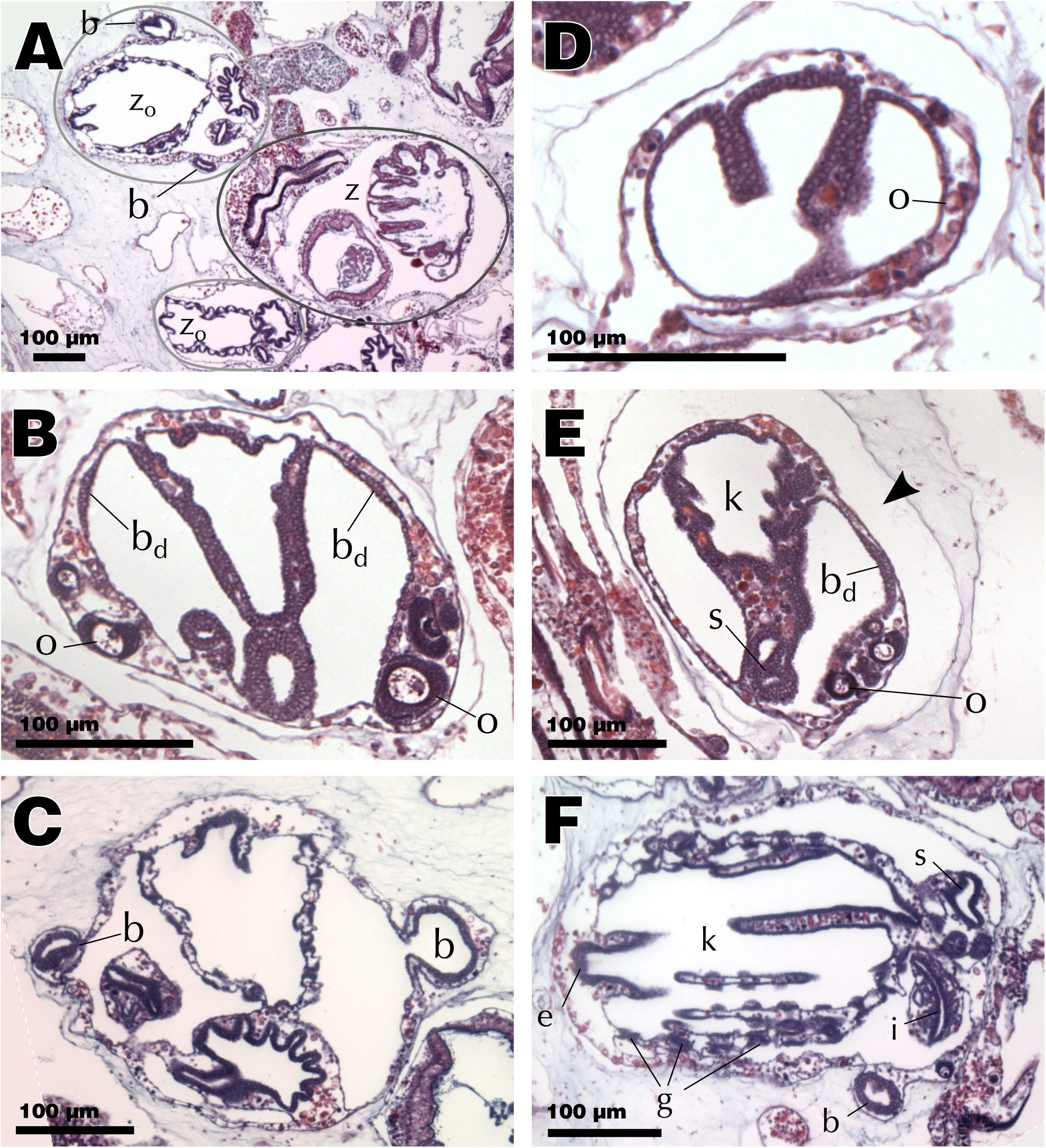
Histology of palleal budding. Abbreviations: zooid (z), first-generation bud (z_0_), second-generation bud (**B**), bud disc (bd), oocyte (o), endostyle (e), pharyngeal basket (k), stigmata (g), intestine (i), stomach (s). All sections are stained with H&E. (**A**) Typical *B*. *leachii* asexual reproduction cycle with three generations co-existing: mature zooid (dark grey), two first-generation buds (light grey) and two budlets visible. (**B**) Bud developing two bud discs and oocytes. (**C**) First-generation bud with right budlet protruding and left budlet fully circular. (**D**) Young bud starting to invaginate. (**E**) Older bud with pharyngeal basket, tentative stigmata, a bud disc and oocytes. Arrowhead points at the interstitial space surrounding the bud. (**F**) Bud on the point of becoming a filter-feeding zooid

The first visible sign of blastogenesis is the appearance of a bud disc, a thickened disk of cells on the ventral side of the atrial epithelium of a first-generation bud, just before the stage where its stigmata become perforated. In healthy colonies, one bud disc usually appears on each side of the forming pharyngeal basket (Fig. 4 B), thus producing two budlets and increasing the overall size of the colony. In well-fed colonies, one of these buds can even divide to give rise to two buds on one side of the zooid, producing one further zooid (Berrill, 1947). The disc cells then proliferate until the bud reaches around 12 cells of diameter (day 1 of the second-generation development), protrudes outside of the first-generation bud through its mantle into the tunic (Fig. 4 C) and curves to form a hollow sphere of cells inside a mantle pouch highly reminiscent of the early stages of blastogenesis (days 2-3). This pouch connects the growing bud with the haemocoel of the mother bud, ensuring that haemolymph and oxygen reach the new bud while the rudiment for the vascular connection to the rest of the colony will be initiated underneath by an outgrowth of the epithelium layer (Berrill, 1947), later producing the radial vessel (Burighel and Brunetti, 1971).

The asexually developing bud continues to increase in size and begins to fold, forming three main compartments, which will form the internal structures of the new adult (days 4-6, Fig. 4 D). It is at that stage that the first signs of germ cells can also be observed (Fig. 4 D). The folds will ultimately join, creating the basic structure for the pharyngeal basket (days 7-8). Once the extending pharyngeal basket has joined the stomach and started to produce its first imperforated stigmata, the stereotypical body plan of the zooid is completed, thus becoming a first-generation bud on which new budlets start to appear (day 9, Fig. 4 E). Following this stage, the first-generation bud will differentiate into a fully functional zooid hence growing in size, finalizing its various organs and vascular system (days 10-22, Fig. 4 F), starting its cardiac activity around day 17.

In total, it takes approximately 22 days at 17 °C to produce a functioning zooid through blastogenesis (Berrill, 1947). Afterwards, the zooid will feed for another 8 days before it gets resorbed through a whole-body apoptosis (< 3 days), in which numerous large macrophagelike cells infiltrate and remove effete cells from the senescent tissues (Brunetti, 2009) and the next generation succeeds.

### *B*. *leachii* whole-body regeneration

The vascular and haematological characteristics of WBR in *B*. *leachii*, are described through comparisons with blastogenesis and intact colonies (Fig. 5). For this analysis, we follow a published classification of WBR into 5 stages (Fig. 1 G; Zondag et al., 2016): Stage 0, injury of the colony; Stage 1, healing of the injury; Stage 2, remodelling of the vascular system; Stage 3, condensation of the ampullae; Stage 4, establishment and development of regeneration niches; Stage 5, fully regenerated zooid.

**Fig. 5.**
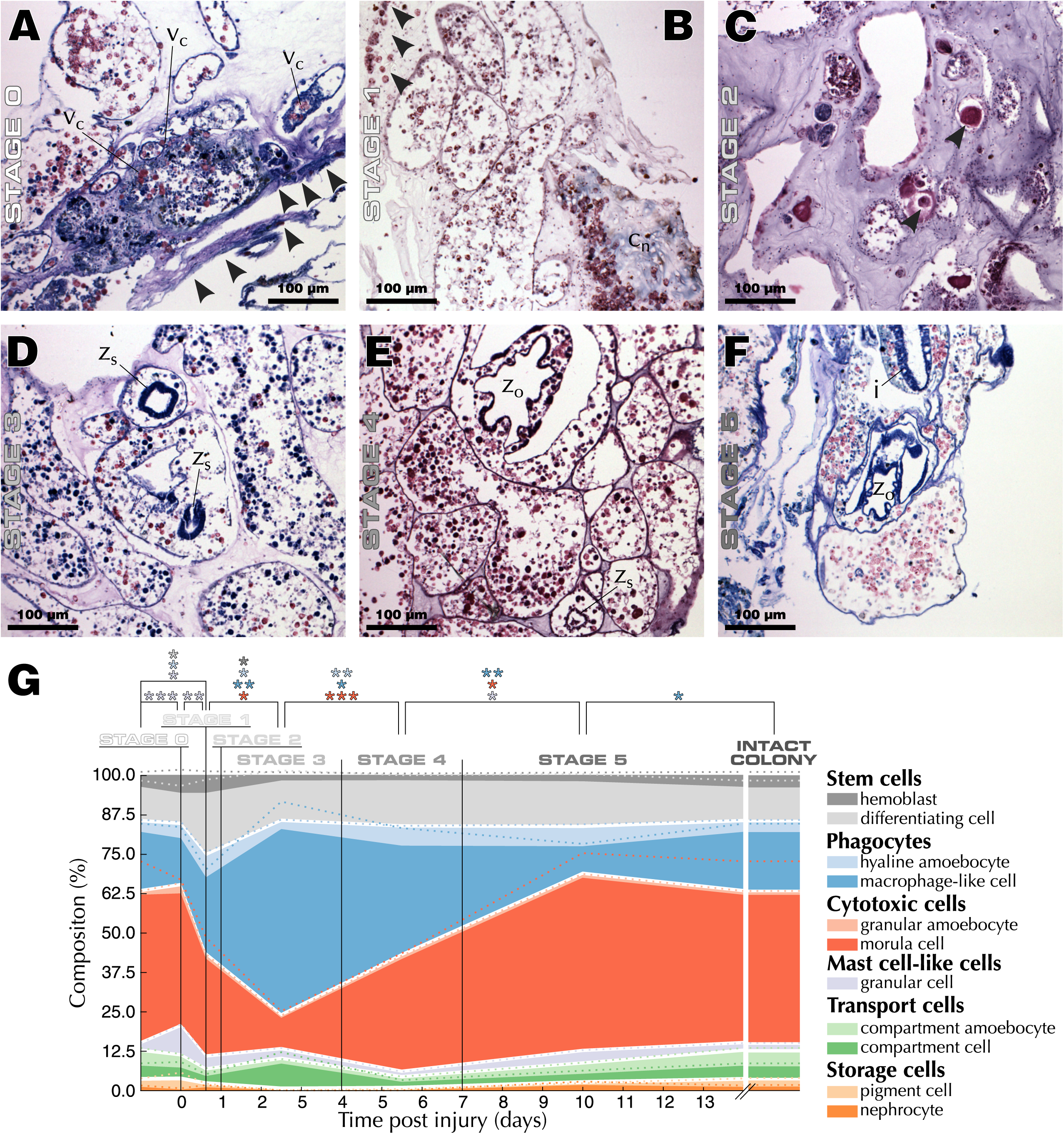
Progression of *B*. *leachii* WBR. Abbreviation: haemolymph clot (v_c_), collagen (c_n_), spheroid bud (z_s_), first-generation bud (z_0_), intestine (i). (**A**) Stage 0: Giemsa stain of injured vascular tissue displaying signs of tunic contraction (arrowheads) and clotting. (**B**) Stage 1: Collagen remodelling and infiltration of cells into the tunic (arrowheads). Methyl blue within MSB dyes collagen blue. (**C**) Stage 2: H&E stained giant cells (arrowheads) throughout the remodelling vasculature. (**D**) Stage 3: Giemsa stained regeneration niches at different stages of development, and increase in macrophage-like cells (see **A** and **F**). (**E**) Stage 4: H&E stain shows competing regeneration niches at the first-generation and spheroid stages. (**F**) Stage 5: Giemsa stain of a fully regenerated zooid with first-generation bud. (**G**) Evolution of haemocyte types within ampullae throughout WBR, overlaid with the corresponding standard error (dotted lines) and stages of WBR (see Fig. 1G). Cell types are indicated in the legend. The color-coded asterisks show statistical significance between the corresponding populations (Student's two-tailed *t-*test, *: *P* < 0.05, **: *P* < 0.01, ***: *P* < 0.001)

#### Stage 0: Injury (0 h)

*B*. *leachii* has a great capacity for preventing haemolymph loss from injuries, with haemolymph loss stopped in less than 30 s by a combination of tissue contraction and clotting (Movie S4). Upon ablation of the zooids, the remaining vascular tissue initially looses a small volume of haemolymph but quickly contracts its peripheral vessels to halt the oozing and starts clotting (Fig. 5 A). Flow will then restart after a pause, the duration of which is seemingly related to either the number of ampullae or the amount of vascular tissue, but was consistently shorter than 10 min. We observed restart of haemolymph flow in as little as two minutes (Movie S4), starting in the portion of the tissue furthest away from the injury in an erratic yet bidirectional fashion, and progressively re-spanning almost the entire vascular system (Movie S4). Although driven only by ampullar contractions, this novel haemolymph flow exhibits a reversal frequency similar to that of intact vessels in a whole colony. As suspected, given the short amount of time available for new cellular differentiation, the composition of this haemolymph is almost identical to the one from intact colonies, with a slight increase in the population of granular cells (Fig. 5 G).

#### Stage 1: Wound healing (15 h)

At this stage of WBR, a dramatic amount of extravascular cellular activity is taking place. The two most apparent aspects are the infiltration of morula cells into the tunic and the remodelling of the tunic matrix (Fig. 5 B). When focusing on circulating haemocytes, we observed an incipient decrease in the population of morula cells, accompanied with an increase in the populations of haemoblasts, hyaline amoebocytes, and granular cells (Fig. 5 G).

#### Stage 2: Ampulla remodelling (24 h)

Concomitant with the observed wound healing process, terminal ampullae start changing their elongated shape to a more spheroid form, contracting, and creating novel tunic vessels (Fig. 5 C; Rinkevich et al., 1995; Rinkevich et al., 2007). One day after injury, previously reported giant cells (Rinkevich et al., 1995, 2007) could be observed throughout the vascular system (Fig. 5 C). In addition, this step exhibits the first signs of the subsequent increase in phagocytic cells (Fig. 5 G).

#### Stage 3: Tissue contraction (2 - 4 d)

By stage 3, the vascular system had fully contracted into a dense network (Rinkevich et al., 1995, 2007) and various regeneration niches were visible within the tissue (Fig. 5 D). Regeneration niches are defined as spherical aggregates of cells within the vascular lumen. Probably the most striking change, at this stage of WBR, was the large increase in phagocytic cells (Fig. 5 D & G), while numbers of both haemoblasts and differentiating cells had returned to pre-injury levels (Fig. 5 G).

#### Stage 4: Regeneration niches (5 - 7 d)

During stage 4, several regeneration niches were present and the foundation of an adult could be observed within one of them (Fig. 5 E). Nonetheless, only one zooid will regenerate and the less developed niches will be resorbed (Rinkevich et al., 1995, 2007). Advanced niches already displayed well developed axial and tissue organizations reminiscent of the early first-generation stage of blastogenesis (Fig. 4 E), including an endostyle as well as an imperforated pharyngeal basket (Fig. 5 E). In comparison to palleal budding, the regenerating buds were lacking both an interstitial space around the niche and visible gametes (compare Fig. 4 E and Fig. 5 E).

At this stage of WBR, haemolymph flow can be observed in novel tunic vessels (Movie S5), and the composition of the haemolymph strives towards the composition observed in intact colonies. Transport, mast-cell like, and storage cells are at their lowest during this period while hyaline amoebocytes and macrophage-like cells remain at a higher proportion than in uninjured colonies (Fig. 5 G).

#### Stage 5: Regenerated zooid (8 - 10 d)

At the conclusion of WBR, a single fully functional zooid is regenerated, displaying the palleal buds expected from an uninjured adult (Fig. 5 F). The haemolymph composition of this new adult is largely similar to that of an intact colony, albeit with a slightly smaller phagocytic cell population (Fig. 5 G).

## Discussion

Here, we carefully dissected the haematological and histological properties of both intact and regenerating *B*. *leachii* colonies. We compared haemolymph flow patterns to published data from related ascidian species and revised previous studies on *B*. *leachii* asexual development In addition, we followed the quantity and location of each type of haemocyte during whole-body regeneration, finding that regeneration progresses through a rapid healing response, an increased activity of haemoblasts, the recruitment of macrophagelike cells and finally their clearing from the haemolymph.

### Synchronization of blood flow in intact and regenerating colonies

Using time-lapse recording, we were able to quantify the complex haemolymph flow patterns observed within a *B. leachi* colony. While the velocity of haemolymph in ascidians has to our knowledge not yet been estimated, the measured average velocity predicts that haemocytes travel an average of 8 mm during one alternation of the flow. In such a small organism (∼2 mm in length), this range could be critical for spreading signalling metabolites and thus potentially coordinating colony responses. The values we obtained for the reversal rates are similar to those measured in other ascidian species, albeit slightly more asymmetric and with a relatively shorter advisceral period (21 to 36 s in *B. leachi*, 30 to 50 s in *Botryllus primigenus* and 45 to 60 s in *Symplegma reptans;* Mukai et al., 1978). We also observed a quick reestablishment of a regular haemolymph flow after injury, and despite the removal of all zooids from the vascular system. This flow is thus sustained solely by synchronized and localized patterns of ampullar contractions (Fig. 2 C & Fig. S3, Movie S4-S5). While this synchronization could solely be a consequence of haemolymph pressure and elastic vascular tissue (Mukai et al., 1978), the absence of any zooid heart within the tunic suggests the existence of a yet unidentified stimulus coordinating this activity.

Similar to vertebrates (Miquerol and Kelly, 2013), *B*. *leachii* heart activity starts before the zooid is fully mature, when neither capable of filter feeding nor of using most of its other organs. Early heart function is consistent with the necessity to circulate oxygen and nutrients throughout the developing organism, even though the parental heart is connected directly to the haemocoel of the bud and would appear to be sufficient for sustaining such circulation. In fact, the pressure exerted by the parental heart is strong enough to impose its direction and reversal rate onto the daughters’ heart (Burighel and Brunetti, 1971).

### Morphological and cellular changes during distinct phases of *B*. *leachii* WBR

We additionally characterised the morphological and haematological modulation of injured colonies throughout WBR. Morula cells are known constituents of the immune system involved in inflammatory responses, haemolymph clotting, homeostasis, and tunic repair (Ballarin et al., 2011, 2001; de Leo, 1992; Menin et al., 2005). The large number of infiltrated morula cells observed during stage 1 of WBR highlights a need for both clearance of decaying or foreign debris and reorganization of the injured vascular system during the first 15 h post dissection. In addition, we measured an increase in the population of differentiating cells (Fig. 5 G), consistent with the need to re-establish homeostasis within the *B*. *leachii* vasculature and to compensate for haemolymph loss. An increased number of mast cell-like cells, a cell type apparently absent from related ascidian species (Cima et al., 2002), was also observed. While their role remains unclear, two hypothesis have been proposed: either a source of nourishment or the immunosurveillance of the alimentary tract (Cima et al., 2002). Their presence during WBR, in the absence of any feeding zooid, rather supports a nutritive role.

At stage 2, we observed the first signs of the subsequent increase in phagocytic cells, potentially as a result of the previously observed differentiation activity (Fig. 5 G) and of the inflammatory response induced by the morula cells (Fig. 5 B). Given the role of these macrophage-like cells in the removal of foreign elements from the tissue (de Leo, 1992), their presence during stage 3 of WBR points towards a recruitment for undertaking the clearing of bacteria, dead cells, and residues of remodelled tunic. The concomitant reduction of haemoblasts to pre-injury levels suggests that the wound healing and remodelling phase has been completed, and that cellular proliferation as well as differentiation have returned to haemopoietic levels within the vascular system. Finally, during stage 4, mast cell-like, transport, and storage cells were observed at their lowest level throughout WBR. This could be a direct consequence of the imposed absence of feeding, leading to a lack of nutrients to be transported or stored. Regenerating *B*. *leachii* colonies would then need to obtain nutrients in an alternative way. Similar to reactivation of hibernating colonies (Burighel et al., 1976) and to the takeover of a new generation of buds in *Botryllus schlosseri* (Lauzon et al., 2002), extraction of reserve nutrients from macrophage-like cells would supplement this need and gradually reduce the number of phagocytic cells, as observed experimentally (Fig. 5G).

When focusing on vascular circulation, the initial rapid interruption in haemolymph flow (Movie S4) could be a consequence of either the reduced density of haemolymph inside the vascular system, or an actively regulated mechanism to reduce the pressure at the sectioned vessels and aiding clotting. Given that haemolymph flow restarts over a relatively short time scale compared to that necessary for substantial haemocytes production and pressure increase, the latter hypothesis appears more likely. This observation thus further supports the existence of an unidentified stimulus systemic to the entire vascular system, potentially relayed directly by the epithelial cells lining the vasculature (Mukai et al., 1978).

In our histological samples, we have observed a detachment of the vascular lining throughout, including in intact colonies (Fig. S4). This suggests that this detachment phenomenon, which has been described as the initial establishment of a regeneration niche (Rinkevich et al., 2007), is either a symptom of vascular remodelling that takes place naturally in healthy colonies but was somehow not prominent in our regenerating vascular tissues, or an artefact of the fixation procedure. Given that we did not consistently detect the reported precursory accumulation of haemocytes at the vascular lining, the latter hypothesis appears more likely, and we suggest that these peripheral cells are morula cells responding to an inflammatory response at the early stages of infiltration.

### Similarities and divergences between blastogenesis and WBR

The two major discrepancies between these two processes were the absence during WBR of an interstitial space around the niche and a lack of visible gametes (Fig. 5 E). As WBR restores both the soma and the germ line (Rinkevich et al., 1995, 2007), the absence of gametes in regenerating niches indicates that they will be created during subsequent cycles of blastogenesis. This process may be an energetically economical developmental approach that avoids the production of gametes in niches that could subsequently be resorbed. Overall, once the vascular tissue has fully contracted, the timing of WBR initially relates closely to that of blastogenesis (day 1-8), but ultimately produces a functional adult much faster. One potential cause for such increased speed is the temperature at which the regenerating colonies were kept (∼19 °C in the present study), which is known to directly correlate with developmental pace (Berrill, 1947, 1941). Temperature alone may not entirely explain such difference in timing however, and one likely cause could be that functional regenerated zooids are smaller than palleal ones, thus having a shortened growth phase which normally spans more than half of blastogenesis. Indeed, such reduction in zooid size was observed in *Botryllus schlosseri* (Voskoboynik et al., 2007).

## Conclusion

Overall, this study underlines the complex interplay of mechanisms required for successful WBR, and complements previous morphological studies (Rinkevich et al., 1995, 2007) by providing a higher temporal resolution. Our histological analysis of WBR supports our recent sequencing approach that identified an initial healing phase followed by a regeneration response (Zondag et al., 2016) and have provided the first detailed account of the haematological properties of *Botrylloides leachii* colonies throughout whole-body regeneration.

## Compliance with Ethical Standards

Conflict of Interest: The authors declare that they have no conflict of interest. Informed consent was obtained from all individual participants included in the study. All applicable international, national, and/or institutional guidelines for the care and use of animals were followed. All permissions required for this study have been obtained.

## Acknowledgements

We would like to thank Francesca Cima (Department of Biology, University of Padova) for pieces of advice, protocols and general help on haemocytes, as well as for proofreading this article; Rueben Pooley (Department of Marine Science, University of Otago) for his help in animal husbandry; Lorryn Fisher (Department of Anatomy, University of Otago) for providing the PCNA antibody; Noel Jhinku (Department of Pathology, University of Otago) for providing the rotifers; Kendall Gadomski (Department of Marine Science, University of Otago) for providing the All-G-Rich powder; Kristian Sveen (Institute for Energy Technology, University of Oslo) for developing the original code of MatPIV; Cynthia A. Brewer (GeoVISTA Center, Pennsylvania State University) for developing ColorBrewer; Fernando Romero Balestra (Department of Cell Signalling, CABIMER), Aude Blanchoud and Anna Jazwinska (Department of Biology, University of Fribourg), for proofreading this article.

The project was supported by a Dean’s Bequest Grant and by the Department of Anatomy [Otago University]; S.B. by the Swiss National Science Foundation (SNSF) [grant number P2ELP3_158873]; and L.Z. by a University of Otago Post-graduate scholarship.

## Electronic supplementary material

### Supplementary figure legends

**Fig. S1 Quantification of haemolymph flow.** (**A**) Raw image overlaying the spatial margin (black) used during the analysis. (**B**) Difference between **A** and its successor frame showing moving haemocytes (white/red) and the added margin (magenta). (**C**) Detection of moving cells thresholded from **B**. (**D**) Average detection of haemolymph movement inside vessels. (**E**) Location of the main vessels thresholded from **D**. (**F**) Identified straight segments of vessels (purple). (**G**) Difference frame used for tracking haemolymph flow. Start (blue) and end (red) of objects’ trajectories are visible. Dashed-delimited area magnified in **I**. (**H**) Difference frame following that depicted in **G**. (**I**) Area magnified from **G**. (**J**) PIV quantification (green) of haemolymph flow inferred using frames **G** and **H**. Dashed-delimited area magnified in **K**. (**K**) Areas magnified from **j**. (**L**) 2D density histogram overlaying PIV measurements (dots), inferred flow velocity (orange), smoothed velocity (brown) and the position of the frames displayed in **A-K** (blue)

**Fig. S2 Quantification of ampullar contractions.** (**A**) Raw frame from the time-lapse recording analysed in Fig. 2B. (**B**) Intensity-based thresholding of **A** used for the detection of ampullae. (**C**) Detection of the ampullae in **A,** color-coded in shades of purple and overlaid with centred circles of corresponding area. (**D**) Clustering of ampullar detections, color-coded in shades of purple. Each circle depicts ampullar detection in some frame of the entire recording. Coloured circles represent detections clustered together, black circles represent spurious detections that were filtered out

**Fig. S3 Ampullar contraction induces haemolymph flow.** (**A**) Raw frame from the time-lapse recording Movie S5. (**B**) Average difference between consecutive frames of the recording. The blue-white-red colour-code depicts contraction in blue and dilatation in red. (**C**) Average absolute second order difference between frames depicting haemolymph flow

**Fig. S4 Vascular lining detaching from the tunic in an adult colony.** High magnification of a Giemsa stained histological section of a terminal ampulla in an adult colony. Arrows point a portion of the vascular lining detaching from the surrounding tunic

**Movie SI** Showcasing the various aspects of haemolymph flow in the vascular system of *B*. *leachii* vascular system using brightfield microscopy. Three different movies (from a large vessel, at a vascular junction and terminal ampullae, respectively) were combined to produce this presentation. The time and scale are specified on the frames of the recording in the top-left and bottom-right corners of the image, respectively. MPEG-4 recording file encoded with the H.264 codec

**Movie S2** Haemolymph flow analysed in Fig. S1. MPEG-4 recording file encoded with the H.264 codec

**Movie S3** The turbulences of haemolymph flow at junctions during flow reversal. The upper panel depicts the measured haemolymph velocity in the various vessel segments, while the lower panel overlays the original recording with the detected vessel segments (blue) and haemolymph flow (red arrows). MPEG-4 recording file encoded with the H.264 codec

**Movie S4** Time-lapse recording of injury-induced WBR. MPEG-4 recording file encoded with the H.264 codec

**Movie S5** Time-lapse recording of WBR, 6 days post injury. The recorded tissue is the one visible in Movie S4. MPEG-4 recording file encoded with the H.264 codec

**Table S1.**
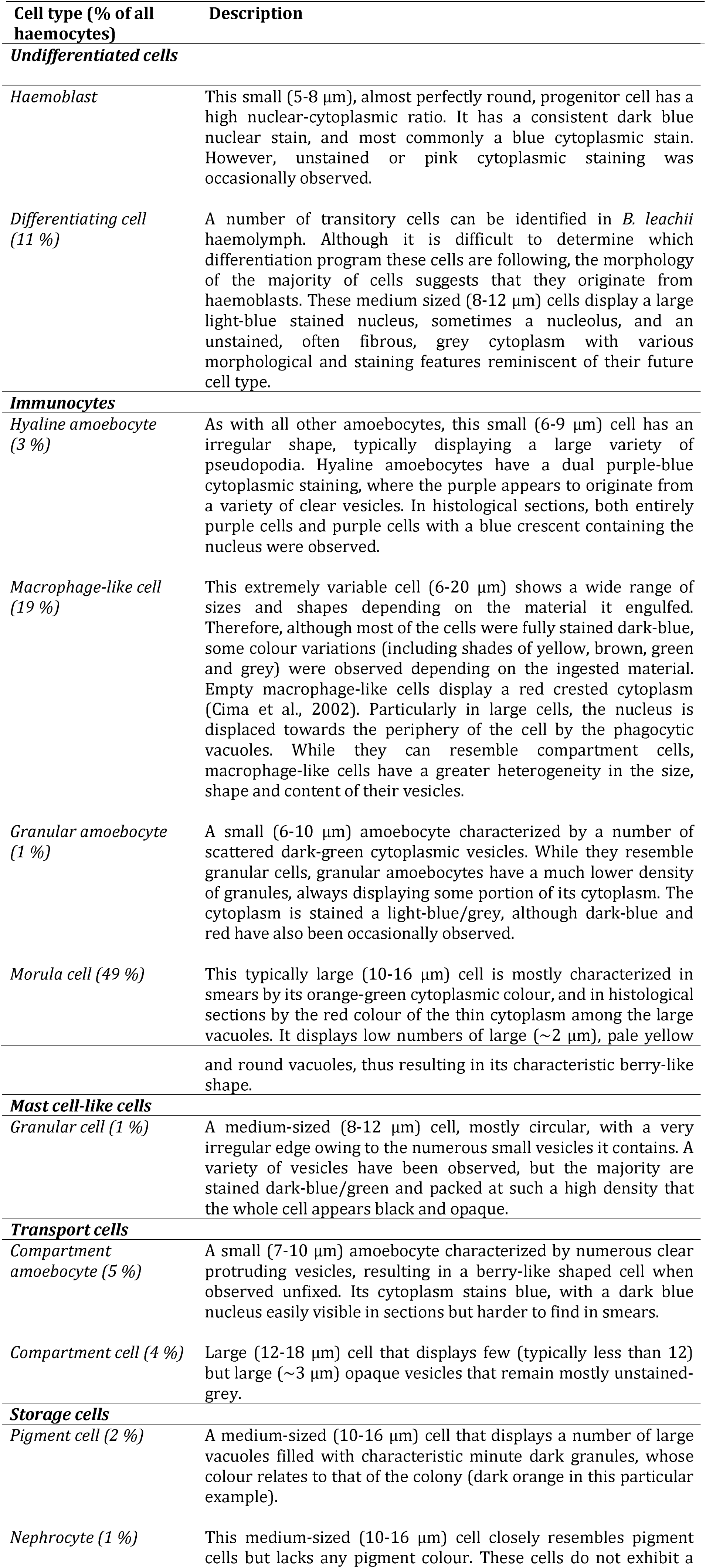
Detailed description of haemolymph cell types present in the *B*. *leachii* vascular

